# Left frontal motor delta oscillations reflect the temporal integration of multimodal speech

**DOI:** 10.1101/2020.11.26.399709

**Authors:** Emmanuel Biau, Benjamin G. Schultz, Thomas C. Gunter, Sonja A. Kotz

## Abstract

During multimodal speech perception, slow delta oscillations (~1 - 3 Hz) in the listener’s brain synchronize with speech signal, likely reflecting signal decomposition at the service of comprehension. In particular, fluctuations imposed onto the speech amplitude envelope by a speaker’s prosody seem to temporally align with articulatory and body gestures, thus providing two complementary sensations to the speech signal’s temporal structure. Further, endogenous delta oscillations in the left motor cortex align with speech and music beat, suggesting a role in the temporal integration of (quasi)-rhythmic stimulations. We propose that delta activity facilitates the temporal alignment of a listener’s oscillatory activity with the prosodic fluctuations in a speaker’s speech during multimodal speech perception. We recorded EEG responses in an audiovisual synchrony detection task while participants watched videos of a speaker. To test the temporal alignment of visual and auditory prosodic features, we filtered the speech signal to remove verbal content. Results confirm (i) that participants accurately detected audiovisual synchrony, and (ii) greater delta power in left frontal motor regions in response to audiovisual asynchrony. The latter effect correlated with behavioural performance, and (iii) decreased delta-beta coupling in the left frontal motor regions when listeners could not accurately integrate visual and auditory prosodies. Together, these findings suggest that endogenous delta oscillations align fluctuating prosodic information conveyed by distinct sensory modalities onto a common temporal organisation in multimodal speech perception.

## INTRODUCTION

Speaker prosody displays perceptible fluctuations in the speech amplitude envelope that allow a listener to segment and parse incoming speech (Ghitza, 2017). While not isochronous, prosody imposes a temporal structure with regular alterations of strong and weak accentuated cues occurring at ~1 - 3 Hz delta rate (Ding et al., 2016; Doelling et al., 2014; Ghitza, 2017; Pell & Davis, 2012). Delta oscillation responses track and align with these prosodic events in auditory cortex to extract the temporal structure of speech (i.e. “neural entrainment”; Giraud & Poeppel, 2012; Keitel et al., 2017; Kosem & van Wassenhove, 2017; Meyer, Sun & Martin, 2019). Beyond segmentation, prosody is present in the visual and auditory domains and may facilitate the listener’s brain activity to synchronize with multimodal information in social interactions (Esteve-Gibert & Guellaï, 2018; Kotz, Ravignani & Fitch, 2018). The term “visual prosody” encompasses communicative gestures (i.e. hand, head, face, and body movements) whose prominent phase temporally coincides with acoustic prosodic anchors such as intonational phrases, pitch accents, and boundary tones (Biau et al., 2016; Chandrasekaran et al., 2009; Munhall et al., 2004; Wagner et al., 2014). For example, the famous cocktail party effect illustrates how listeners rely on temporal alignment between gestures and sounds to improve speech perception (Cherry, 1953; Obermeier, Dolk & Gunter, 2012; Sumby & Pollack, 1954). Together these facts raise the following question: How does the brain integrate multiple dynamic visual and auditory prosody streams to facilitate multimodal speech perception?

The present study investigated how delta oscillations mark the temporal integration of audiovisual prosody in speech. We refer to temporal integration as the mechanism that integrates visual and auditory prosody, leading to the improved perception of their temporal representation in speech. Delta activity in the motor cortex has been associated with the temporal integration of rhythmic stimuli including speech, as its phase aligns with the onsets of predictable events (Morillon et al., 2019; Morillon & Schroeder, 2015; Saleh et al., 2010). Keitel et al. (2018) showed that left motor delta activity tracked temporally predictable slow phrasal features in auditory sentences and predicted speech comprehension. This suggests that this region integrates perceptually relevant regularities in the signal to facilitate comprehension. The authors also found delta-beta cross-frequency coupling in the left motor region, in line with previous research showing that motor beta oscillations also respond to the temporal integration of rhythmic auditory tones or visual cues stimulations (Fujioka, Ross & Trainor, 2015; Saleh et al., 2010). These findings led to the hypothesis that motor delta oscillations are involved in the temporal integration of speech by mediating top-down control through cross-frequency coupling with beta activity (Arnal, 2012; Arnal, Doelling & Poeppel, 2015; Morillon and Baillet, 2017). In other words, delta activity could reflect how the brain gathers multiple temporal representations of input across modalities in the left motor cortex, and generates predictions to improve signal processing. Finally, the left motor cortex including the left inferior frontal gyrus is involved in gestures and speech integration, making it a critical candidate of the present study (Biau et al., 2016; Park et al., 2016; Zhao et al., 2018).

We propose that visual and auditory prosodic cues encoded in the visual and auditory sensory cortices respectively, provide two representations of the speech signal that are integrated in the left motor in speech perception. Such crossmodal temporal integration is reflected by delta responses in this region. To test this hypothesis, we manipulated the temporal alignment of filtered speech with corresponding whole body or masked head movements. Participants performed an audiovisual synchrony detection task by attending to videos of a single speaker engaged in a conversation, while we recorded their electroencephalogram (EEG). First, we tested whether listeners relate the respective temporal structures of visual and auditory prosodic features in multimodal speech. Second, we explored delta oscillations in response to audiovisual asynchrony in multimodal stimuli. Third, we tested whether delta-beta coupling in the left motor cortex predicted multimodal synchrony detection in speech perception.

## METHODS

### Participants

Twenty-six native Dutch speakers (mean age = 22.24, SD = 4.24; 15 females) were recruited at Maastricht University and received €10 for participating in the experiment after giving informed consent. All participants were right-handed and had normal or corrected-to-normal vision and hearing. The protocol of the study was approved by the Research Ethical Committee of Maastricht University. Data from three participants were removed from the final analysis due to technical problems.

### Stimuli

Short videos were extracted from a longer video recording used in a previous study (Gunter & Weinbrenner, 2017). The videos depicted a female actor and an experimenter (both German native speakers) engaged in a question-answer conversation. The actor sat on a chair, moved freely, and was visible from her knees up to the top of her head. Relevant segments containing the actor’ answers separate from the experimenter were selected to create the current stimulus set (N = 54). Each of the 54 segments was 10 seconds long (600 frames at 60 frames per second; FPS). The audio track was extracted to be low-pass filtered with Hann band windowing procedure (from 0 Hz to 400 Hz; 20 Hz smoothing) using Praat (Boersma & Weenink, 2015). In doing so, we altered speech intelligibility removing verbal content while keeping the prosodic contour of the signal. Peak frequencies were extracted from the audio and video files through Fourier transformations that calculated the frequency at which the peak amplitude occurred within a range of 0.5Hz to 8Hz. For videos, the average magnitude of grayscale pixel changes between consecutive frames was used to determine the frequency of movement and gesture (see Table 1).

**Table 1.** Summary statistics of peak frequencies obtained for video and audio signals using a Fourier transformation (*Features remaining identical across the no mask and head mask conditions).

| Row Labels | Mean Peak Freq. | SD Peak Freq. | Min. Peak Freq. | Max Peak Freq. |
| --- | --- | --- | --- | --- |
| **Audio\*** | 2.74 | 1.44 | 0.86 | 5.86 |
| **Body\*** | 3.37 | 1.25 | 0.86 | 6.13 |
| **Head-mask Face** | 2.27 | 1.53 | 0.86 | 6.13 |
| **Head-mask Full** | 3.10 | 1.45 | 0.86 | 6.13 |
| **No-mask Face** | 3.59 | 0.91 | 1.00 | 3.99 |
| **No-mask Full** | 3.65 | 1.02 | 0.86 | 6.13 |

We applied two visual manipulations to each of the 54 speech segments: (1) The presence or absence of a visual mask (no mask, head-mask), and (2) the original temporal alignment of the audiovisual information or a temporal shift of the audio signal relative to the video onset (synchronous, asynchronous). In the no-mask condition, the speaker’s body and face were fully visible. In the head-mask condition, the head of the speaker was blurred to degrade visual prosody conveyed by the speaker’s lips. The mask was created by applying a low-pass Gaussian filter on the upper third of the original video containing the speaker’s face, attenuating a high frequency signal. This manipulation removed fine-grained facial expressions from the video while slow gestures remained intact (see Figure 1). In the synchronous condition, the original temporal alignment between visual and auditory onsets was intact. To create an asynchronous condition, we inserted a delay between the visual and auditory onsets by shifting the sound onset by +400 ms relative to the video onset (i.e., 24 frames). This manipulation maintained the natural order of visual information preceding auditory information. A 400ms lag was used to ensure that the delay was long enough to detect audiovisual asynchrony; it was also based on the time-window of multisensory integration established in previous studies (Biau et al., 2016; Biau & Soto-Faraco, 2013; Jessen & Kotz, 2015; Obermeier & Gunter, 2014). Further, a central white fixation cross was displayed in each video to allow participants to focus their gaze on a central cue while attending audiovisual stimuli. Altogether, this created four conditions: Head-Mask Synchronous (HMS), Head-Mask Asynchronous (HMA), No-Mask Synchronous (NMS), and No-Mask Asynchronous (NMA) (see Figure 1A). 18 additional video clips, in which the central white fixation-cross turned red, were used as fillers, counterbalanced across conditions (colour change onset jittered between 5 and 9 seconds after the video onset; ~ 8 % of total stimuli, not included in the final data analysis). We used the fillers in a memory test to focus the participants’ attention on the videos during the experiment. Finally, audio files were recombined with their corresponding video files for each condition. Videos were edited using Adobe Premiere Pro CS3 and exported using the following parameters: Pixel resolution 1920 × 1080, 60 FPS compressor Indeo video 5.10, AVI format, audio sample rate 48 kHz, 16 bits Mono.

**Figure 1.**
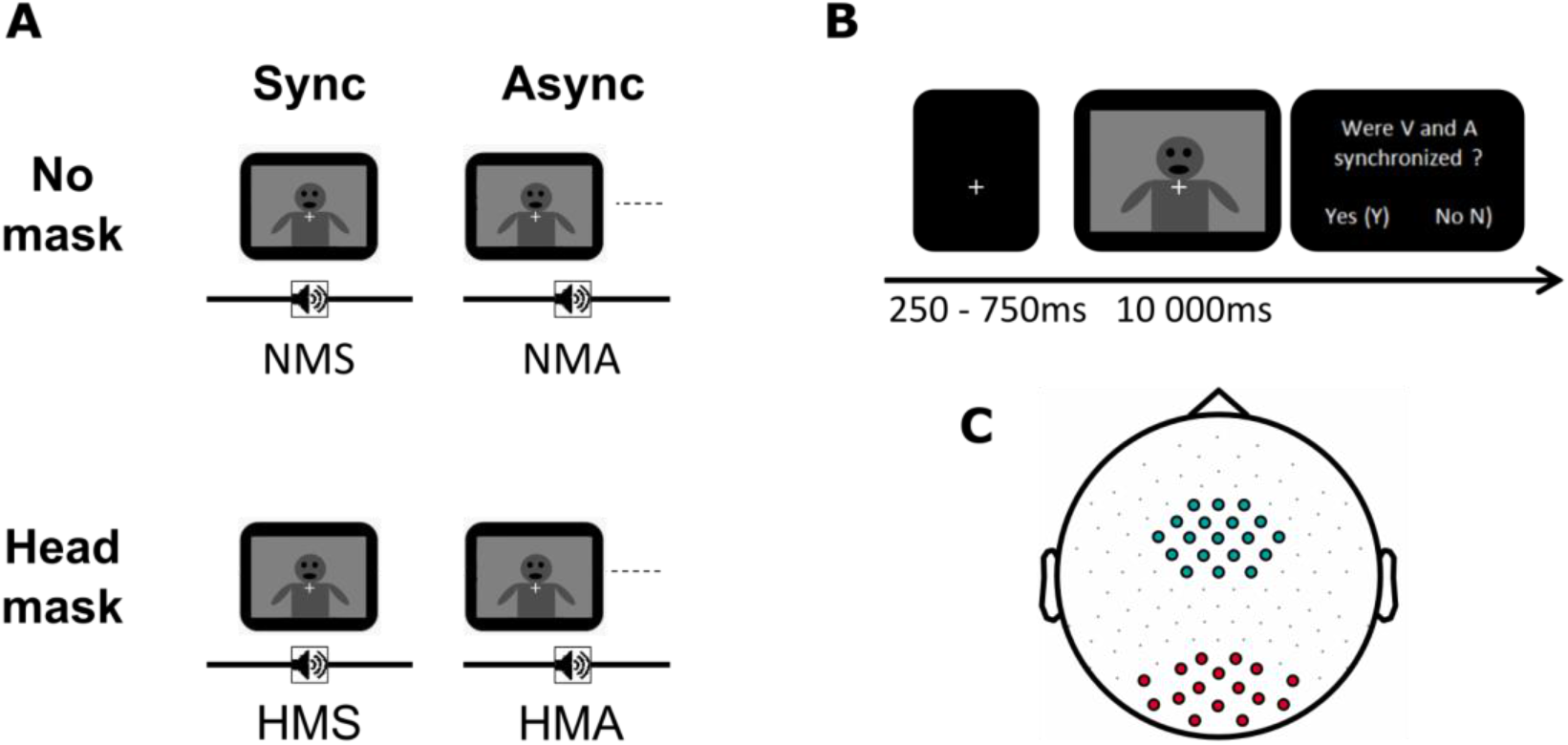
Experimental procedure of the audiovisual synchrony detection task. (A) For each item, the audio signal was the same across all four versions. Visual information was manipulated for the factor mask (no-mask or head-mask); audiovisual stimuli were temporally aligned in the synchronous conditions (NMS, HMS), and the audio signal was temporally delayed (400ms) in the asynchronous conditions (NMA, HMA). (B) Example of one trial timeline. (C) Distribution of the electrodes covering the motor region of interest (ROI; blue circles) and the control region of non-interest in the visual area (RONI; red circles). N.B: The images in A and B have been modified for anonymity purpose here.

### Apparatus

The audio files were presented through EEG-compatible air tubes (ER3C Tubal Insert Earphones, Etymotic Research). Videos were presented on a 27 inch Iiyama G-MASTER (GB2760HSU-B1) TN display with a 1ms response time, a refresh rate of 144Hz, and a native resolution of 1920 x 1080 pixels connected to the stimulus presentation computer (Intel i7-6700 CPU @ 3.40 GHz, 32 GB, running 64-bit Windows 7, NVIDIA GeForce FTX 1080 GTX GPU). Stimuli were presented using a custom MATLAB script (MATLAB and Statistics Toolbox Release 2015b, The MathWorks, Inc., Natick, Massachusetts, United States) that called VideoLAN Client (VLC; VideoLAN Client, 2017; http://www.videolan.org/) to play the videos. EEG data were collected using BrainVision Recorder (Brain Products, GmbH, 2017) software on an Intel Xeon E5-1650 PC (3.5 GHz, 32GB RAM) running Windows 7. Video onsets were synchronized to EEG data using the Schultz Cigarette Burn Toolbox (Schultz, Biau, & Kotz, 2020).

### Procedure

Participants were seated approximately 60 cm apart from the monitor in a sound attenuated booth while videos were displayed on a computer screen. Participants watched 234 videos organised in nine blocks of 26 randomised trials (i.e., 6 stimuli per condition + 2 fillers). The task was a two-alternative forced choice synchrony detection task (Figure 1B). Participants attended both the audio and video stimuli. Each trial began with a central white fixation cross (jittered duration 500 +/− 250 ms), followed by the stimulus. After the video ended, participants decided whether the audio and the video signals were synchronous or asynchronous by pressing the “1” or “2” key on the keyboard without time pressure (counterbalanced across participants). Additionally, participants were asked to count internally the number of times they observed a red cross in a video clip, and reported it at the end of the experiment. Filler trials were not included in behavioural and EEG analyses but the total number of reported red crosses served to check that attention was maintained throughout the experiment. Before the experiment, participants received five practice trials where they were presented with one example of each condition to ensure they understood the instructions. At the end of the experiment, participants were asked if they could identify the speaker’s language and to report it.

### EEG recording and preprocessing

Electrophysiological data were recorded at 1000 Hz with 128 active electrodes (ActiCap, Brain Vision Recorder, Brain Products) according to the 10-20 international standard, and impedances were kept below 10 kΩ. The ground electrode was located at AFz, and the reference electrode was placed at the right mastoid (TP10).

Offline EEG preprocessing: EEG data were preprocessed offline using Fieldtrip (Oostenveld et al., 2011) and SPM8 toolboxes (Wellcome Trust Centre for Neuroimaging). Continuous EEG signals were bandpass filtered between 1Hz and 100 Hz and bandstop filtered (48-52 Hz and 98-102 Hz) to remove line noise at 50 and 100 Hz. Data were epoched from 1000 ms before stimulus onset to 11000 ms after stimulus onset. Trials and channels with artefacts were excluded by visual inspection before applying an independent component analysis (ICA) to remove components related to ocular artefacts. Excluded channels were then interpolated using the method of triangulation of nearest. After re-referencing the data to an average reference, the remaining trials with artefacts were manually rejected by a final visual inspection (on average, 13.57 ± 8.32 trials across conditions per participant).

### EEG data analyses at the scalp level

For each participant, time-frequency representations (TFRs) were computed using a Morlet wavelet (width: 5 cycles) from 1 to 40 Hz (1 Hz step), with 20 ms time steps. Power in the hit trials (i.e. correct synchrony detection in synchronous conditions and correct asynchrony detection in the asynchronous condition) was calculated first and then averaged across trials in the four conditions. The power was normalised relative to a pre-stimulus baseline (−700 to −200 ms with respect to stimulus onset) to determine increases or decreases of power dependent on the conditions. As entrainment necessitates several cycles from recurrent stimulations to build up (Doelling et al. 2014; Thut et al., 2011; Zoefel et al. 2018) and the slower frequency in our band of interest was 2Hz (corresponding to a period of 500 ms), we defined a time window of interest from + 3 to + 9 seconds after stimulus onset. This time window ensured that neural activity sufficiently entrained to the temporal structure of the stimuli, and that the responses evoked by the stimulus onset-offsets did not influence the results. In the identified regions of interest and non-interest (see Results section), normalised mean power across pool electrodes in the 2-3 Hz frequency band was computed for the four conditions and exported for further statistical analyses.

### EEG data analyses at the source level

#### Source localisation

We used the Montreal Neurological Institute (MNI) MRI template and a template volume conduction model from Fieldtrip. The 128 electrode positions on the volunteer’s head were defined by using a Polhemus FASTRAK device (Colchester), recorded with the Brainstorm toolbox implemented in MATLAB (Tadel et al., 2011), and realigned to the template head model using Fieldtrip. The template volume conduction model and the electrode template were used to prepare the source models. Leadfields were computed based on scalp potentials and source activity was reconstructed applying a linearly constrained minimum variance (LCMV) beamforming approach implemented in Fieldtrip (van Veen et al., 1997; Wang et al., 2018). Source analyses were run on potential data (i.e., average referenced) and time-series data were reconstructed on 2020 virtual electrodes for each participant. Time-frequency analysis was computed at each of 2020 virtual sources with the exact same approach to scalp level analyses. The maximum voxel activation regions were defined by using the automated anatomical labelling atlas (AAL).

#### Phase-amplitude coupling (PAC) between delta and beta oscillations

We applied a modulation index (MI) analysis in the time-window of interest to quantify delta-beta PAC in the significant cluster revealed by source localisation in the NMA-NMS contrast (Tort et al., 2010). First, the power spectrum (1 - 30 Hz) was estimated across all grids of the significant cluster and trials by applying a 1/f correction time-frequency decomposition method with wavelet for each participant (Griffith et al., 2019). The most prominent power spectrum peaks in the delta and beta bands were then extracted and saved as the individual delta and beta peaks. Across participants, the mean delta peak was at 2.38 Hz and the mean beta peak was at 24.16 Hz. The number of trials were balanced by identifying the condition with the smallest number of incorrect trials, and taking 80 percent of the smaller sample for all the conditions (NMA_hit_, NMA_miss_, NMS_hit_ and NMs_miss_; HMA_hit_, HMA_miss_, HMS_hit_ and HMs_miss_). Subsampled trials were concatenated and the operation was repeated for 50 iterations in each condition (Keitel et al., 2018). The grids of interest resulted from the significant cluster identified in the left frontal motor cortex by the source localisation analysis (contrast NMA-NMS; number of significant grids = 92). Phase and power were derived from Hilbert-transformed time series and filtered around the delta peak (± 0.5 Hz) and beta peak (± 5 Hz) based on the frequency window. For each trial and grid source, beta power was binned into 12 equidistant bins of 30° according to the delta phase. The MI was computed by comparing the observed distribution to a uniform distribution for each trial and grid. The PAC was then averaged across the left frontal motor grids and 50 iterations in each condition. Finally, we investigated whether the delta-beta coupling was specifically localised in the region of interest, identified by the source localisation analysis (i.e., left frontal motor area). We compared the delta-beta PAC between masks (no-mask, head-mask) based on the results from the cluster of interest. In contrast to the PAC analysis in the region of interest, the difference of trial numbers between conditions was balanced by taking 80 percent of the smaller sample between all the conditions. Subsampled trials were concatenated and the operation was repeated for 40 iterations (to circumvent computational resource limits reached by concatenated epoch lengths). The delta-beta PAC was then averaged across all iterations at each grid (n = 2020) and conditions across participants.

### Experimental design and statistical analysis

#### Audiovisual synchrony detection task

The experiment used a full within-subject design. The effect of asynchrony and its interaction with the head-mask in audiovisual integration was assessed by means of *d’* sensitivity index and reaction times for correct trials. The correct responses (Hits and correct rejections in the synchronous conditions) and errors (misses and false alarms FA in the asynchronous conditions) as well as the reaction times of the hits (comprised between mean reaction times ± two standard deviations range), were computed in each condition for each participant. Then, the d’ scores for synchrony detection in the no-mask and head-mask contrasts were calculated for each participant as follows: *d’* = Z (Hit_rate_) – Z (FA_rate_). The *d’* index allows taking into account response bias by comparing hits and false alarms to assess whether participants actually discriminated synchrony and asynchrony. Additionally, the decision criterion *c* was computed as follows: *c* = 0.5 x (Hit_rate_ – FA_rate_) / 2 to determine the decision shift between no-mask and head-mask contrasts. The effects of masking the speaker’s face and audiovisual synchrony on reaction times were assessed using two-way repeated-measure ANOVAs with the factors mask (no-mask, head-mask), synchrony (synchronous, asynchronous), and the interaction between mask and synchrony, using SPSS (IBM Corp. Released 2015. IBM SPSS Statistics for Windows, Version 23.0. Armonk, NY: IBM Corp.). In the case of significant interactions, *post-hoc t*-tests were Bonferroni-corrected. To test for the modulation of sensitivity to synchrony depending on speaker’s face information, the *d’* and *c* criterion in no-mask and head-mask contrasts were individually tested against zero by means of one-sample *t*-tests. Further, the difference of *d’* between the no-mask and head-mask contrasts was assessed applying a paired-samples *t*-test and the effect size was defined using Cohen’s *d*.

#### EEG data at the scalp level

EEG data of correct trials at the scalp and source levels were statistically analysed. The differences of mean power between two contrasts (NMA-NMS and HMA-HMS) at the electrode level were statistically assessed by applying dependent *t*-tests using Monte-Carlo cluster-based permutation tests (Maris & Oostenveld, 2007) with an alpha cluster-forming threshold set at 0.05, three minimum neighbour channels, 5000 iterations, and cluster selection based on maximum size. Cluster-based permutation statistics were applied for the time window of interest in the delta 2-3 Hz band across all the electrodes. To address whether centro-frontal delta oscillations responses reflect temporal speech analysis rather than pure signal processing via neural entrainment, we performed the same tests on the theta band (4 - 8 Hz), which plays a role in the integration of the syllabic structure of speech (Giraud & Poeppel, 2012). We expected to find modulations of delta but not theta oscillations for audiovisual asynchrony in the region of interest if motor delta responses reflect temporal integration. In the identified regions of interest and non-interest (see Results section), normalised mean power across pool electrodes in the 2-3 Hz delta and 4-8 Hz theta frequency bands was computed for the four conditions and exported. Statistical differences of power in relevant contrasts were assessed by means of two-way repeated-measure ANOVAs.

#### EEG data source localisation

Differences in delta power for the two contrasts (no-mask: NMA-NMS and head-mask: HMA-HMS) were assessed by applying dependent *t*-tests using Monte-Carlo cluster-based permutation tests as performed for the scalp level analyses. For visualisation of the source localisation results, the power differences in the two contrasts were grand-averaged across participants, and the grand average power differences were interpolated to the MNI MRI template for visualization. Only voxels surpassing the statistical significance threshold are depicted in both contrasts (significant *t*-values at alpha = 0.05, multiple comparison cluster-corrected).

#### Delta-beta PAC

First, statistical differences of mean PAC across conditions in the region of interest were assessed applying a three-way repeated-measure ANOVA with the factors mask (no-mask and head-mask), synchrony (synchronous and asynchronous), and correctness (correct and incorrect trials). Second, statistical differences of whole brain delta-beta PAC was assessed by applying dependent *t*-tests using Monte-Carlo cluster-based permutation tests as described above (whereas *t*-tests were one-tailed here as we had a strong hypothesis about delta-beta PAC modulation directionality based on results at region of interest level).

#### Correlations between synchrony detection performance and delta oscillations in the identified left motor cluster

We addressed whether delta responses in the left motor area correlated with audiovisual synchrony perception. We performed Pearson correlations between the difference of mean power in the 2-3 Hz band at source level, and the difference of correct response rates within contrast. For each participant, we computed the 2-3 Hz power at the grids sources from the significant cluster established in the NMA-NMS contrast source analysis (all significant grids were situated in the left central and frontal gyrus areas; n = 92). Power was averaged across the 92 grids in the four conditions separately (NMS, NMA, HMS and HMA), and we calculated the mean difference separately in the no-mask (NMA-NMS) and head-mask (HMA-HMS) contrasts to obtain two delta power values per participant. Similarly, the difference of correct response rates was calculated in the no-mask and head-mask contrasts, resulting in two behaviour values per participant. The statistical relationship between behaviour and delta power was assessed applying Pearson’s correlation tests.

## RESULTS

Participants reported 18.26 ± SD = 1.51 red crosses (out of 18) at the end of the experiment. Additionally, they correctly identified the speaker’s native language (they all responded “German”), although they could not report any semantic content. These results confirmed that participants correctly paid attention to both the audio and video signals.

### Listeners successfully temporally aligned visual and auditory prosodic features to achieve multimodal speech integration

D’ scores are reported in Figure 2A. Two independent one-sample *t*-tests revealed that the mean *d’* in the no-mask (NMS and NMA) and head-mask contrasts (HMS and HMA) were significantly greater than *μ* = zero (no-mask: *t*(1,22) = 10.25; *p* < 0.001; *Cohen’s d* = 3.04; head-mask: *t*(1,22) = 8.07; *p* < 0.001; *Cohen’s d* = 2.38). A paired-samples *t*-test performed on *d’* for no-mask and head-mask contrasts confirmed that participants detected synchrony better in the no-mask contrast than in the head-mask contrast (*t*(1,22) = 6.96; *p* = < 0.001, two-tailed; *Cohen’s d* = 1.46). Finally, two independent one-sample *t*-tests revealed that the mean *c* criterion in the no-mask contrast was not significantly different from zero (0.11 ± 0.40; *t*(1,22) = 1.32; *p* = 0.1; *Cohen’s d* = 0.39), whereas it was significantly below zero in the head-mask contrast (−0.53 ± 0.28; *t*(1,22) = −9.01; *p* < 0.001; *Cohen’s d* = 2.68). These results established that when the speaker’s face was head-masked, participants were significantly biased to respond “synchrony” when the stimulus was asynchronous than in the no-mask contrast (i.e., liberal guessing). A two-way repeated-measure ANOVA revealed a significant main effect of mask on reaction times (*F*(1, 22) = 16.50, *p* < 0.01; *η_p_^2^* = 0.43). No significant effect of synchrony (*F*(1, 22) = 0.67, *p* = 0.42; *η_p_^2^* = 0.03) or interaction between mask and synchrony was found (*F*(1, 22) = 2.32, *p* = 0.14; *η_p_^2^* = 0.1). These results showed that accurate responses were faster when the face of the speaker was not masked compared to head-masked, in line with greater difficulty in integrating video and audio signals together when visual information was degraded (Figure 2B).

**Figure 2.**
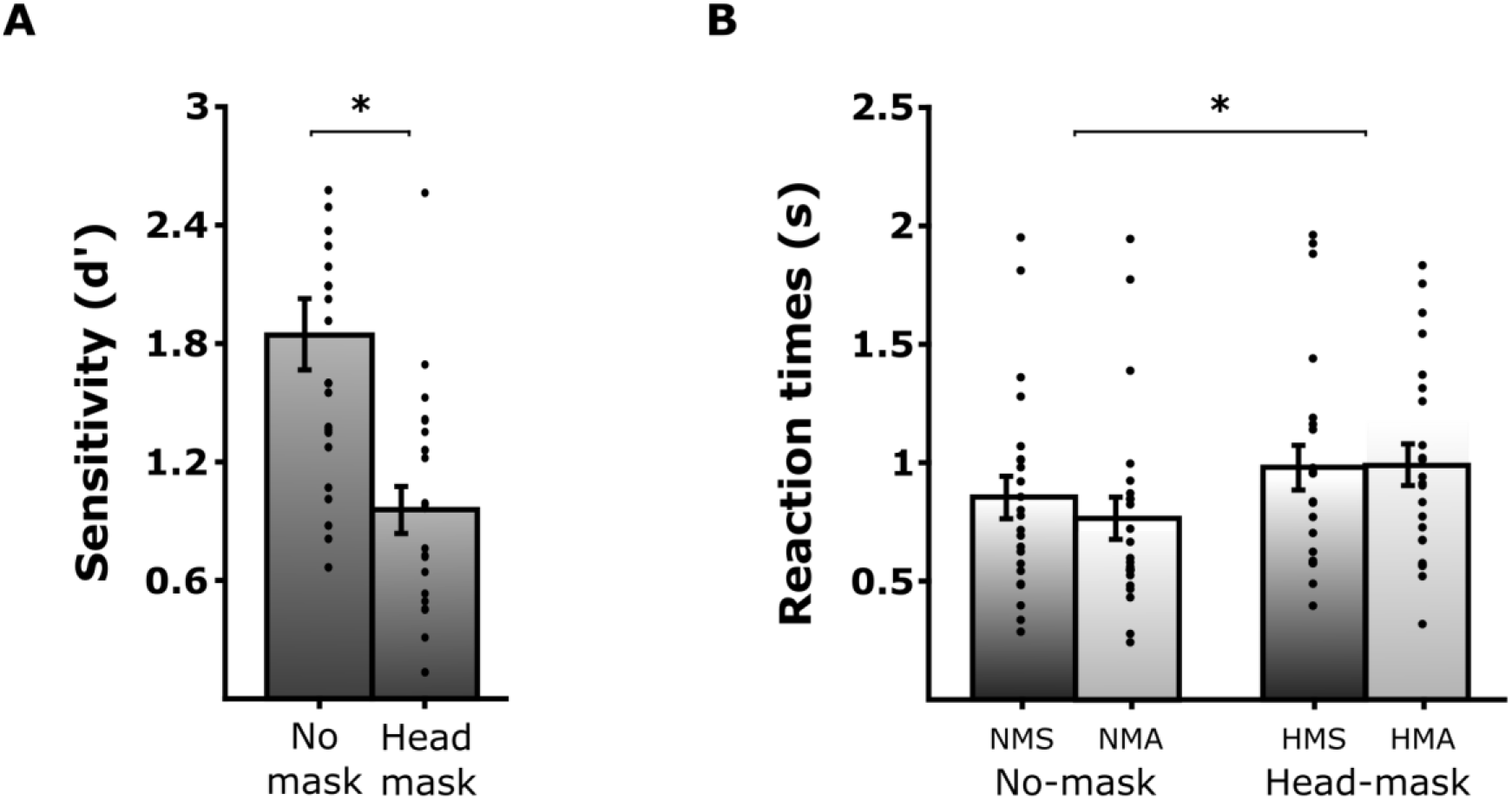
Behavioural performances in the synchrony detection task. (A) Average *d’* scores and reaction times of hits across conditions (± standard error of the mean; grey dots represent individual averages; n = 23). Significant contrasts are marked by stars (*p* < 0.05).

Together, the behavioural performance supports the hypothesis that participants integrated the temporal structure of slow prosodic features in integrating visual and auditory information during multimodal speech perception. Further, when visual information carried by the speaker’s face was degraded with a head-mask, the sensitivity to audiovisual synchrony decreased. This suggests that successful multimodal integration of speech requires visual information of the head and face.

### Delta oscillations in the left frontal-motor cortex reflect temporal integration of audio and visual prosody and shape multimodal speech perception

We then addressed whether delta oscillations in the left motor cortex relate to multimodal temporal integration, and whether responses depend on the amount of visual information available. First, a cluster-based permutation tests revealed a significant increase in delta power (2-3 Hz) in response to the audiovisual asynchrony when the speaker’s face was visible (no-mask: NMA-NMS) but not when it was masked (head-mask: HMA-HMS) (NMA-NMS: *p* < 0.001, cluster statistic = 117.23; HMA-HMS: zero positive cluster statistic; multiple comparisons are cluster-corrected). No significant negative clusters were found in both contrasts. Importantly, the topography of the significant delta cluster in the no-mask contrast showed a main fronto-central response when video and audio signals were asynchronous, in line with the expected source localization of delta in the motor region (Figure 3B; Puzzo et al., 2010; Stegemöller et al., 2017). To assess the potential interaction of visual information and audiovisual synchrony perception in this motor region of interest, we defined a set of electrodes as the region of interest (ROI) representative of the delta response topography: F1, Fz, F2, FFC3h, FFC1h, FFC2h, FFC4h, FC3, FC1, FCz, FC2, FC4, FCC3h, FCC1h, FCC2h, fCC4h, C1, Cz and C2 (Figure 1C). The mean delta power across the electrodes of the ROI was computed in the four conditions separately, and confirmed an increase of induced delta activity compared to the pre-stimulus baseline (NMS: 0.64 ± 0.17; NMA: 0.74 ± 0.15; HMS: 0.70 ± 0.16 and HMA: 0.68 ± 0.20; see Figure 3A and 3C). A two-way repeated-measure ANOVA revealed a significant interaction between the factors mask and synchrony for delta power (*F*(1, 22) =5.78, *p* = 0.03; *η_p_^2^* = 0.21). Bonferroni-corrected pairwise comparisons showed that in the no-mask contrast, delta power was significantly greater in the asynchronous (NMA) than synchronous (NMS) condition (*p* < 0.001). In contrast, asynchrony did not affect delta power responses in the head mask contrast (*p* = 0.52). Second, when video and audio signals were presented in asynchrony, delta power was significantly greater when the speaker’s face was not masked as compared to head-masked (NMA vs. HMA; *p* = 0.038). When the video and audio signals were synchronous, the presence of the mask on the speaker’s face did not significantly modulate the delta power (NMS vs. HMS; *p* = 0.11).

**Figure 3.**
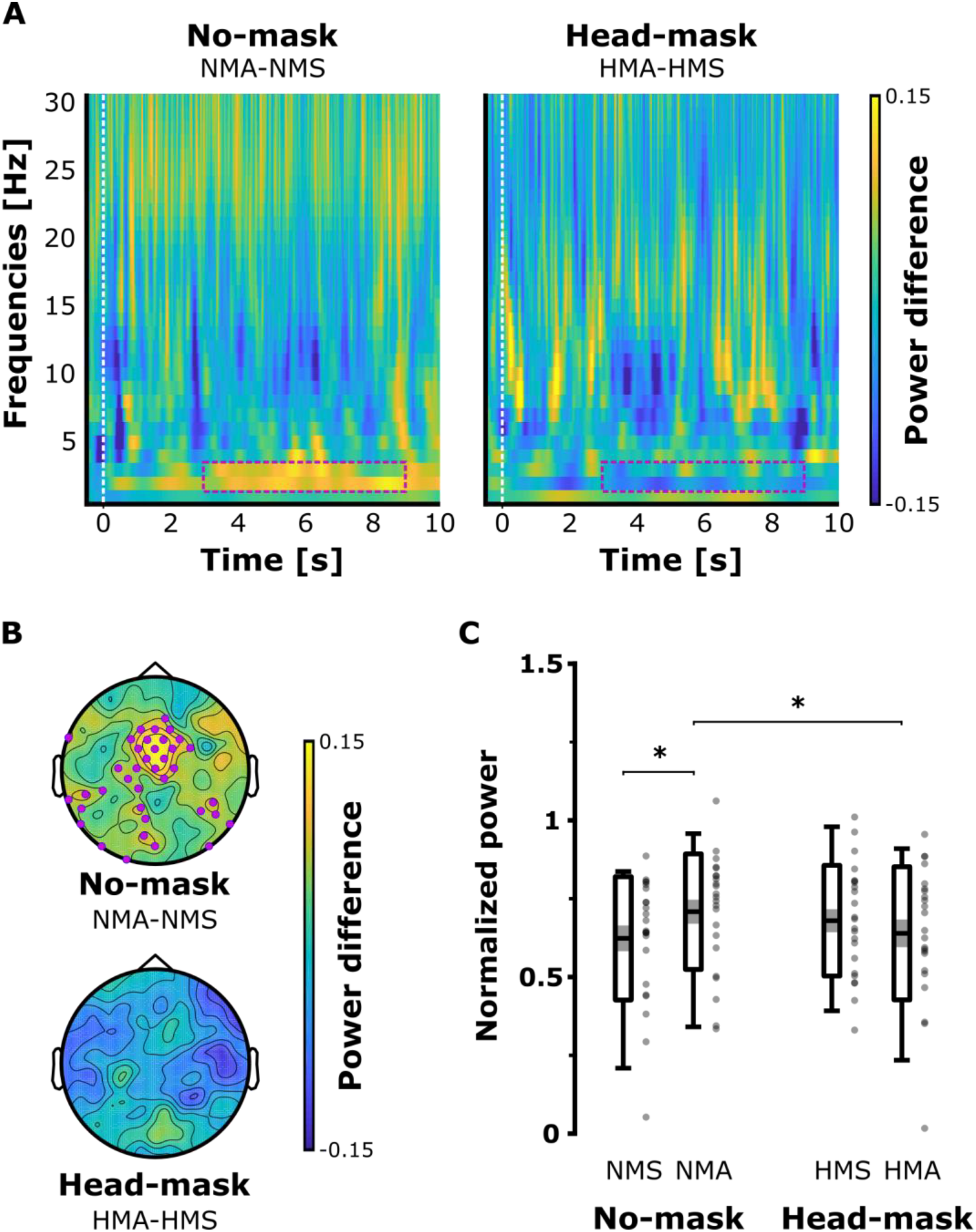
Delta responses to audiovisual asynchrony at the scalp level. (A) Time-frequency spectra of the mean power differences in the motor ROI between asynchronous and synchronous conditions in the no-mask (NMA-NMS; left) and head-mask (HMA-HMS; right) contrasts. The white dashed lines correspond to the onset of the video and the window of interest is marked by the pink dashed rectangles. (B) Topographical distribution of the difference of 2-3 Hz delta power in the time-window of interest, in the no-mask (NMA-NMS; top) and head-mask (HMA-HMS; bottom) contrasts. The pink dots display electrodes with significant *t*-values (alpha threshold = 0.05). (C) Delta power across the electrodes of interest in the four conditions (2-3 Hz band). Significant contrasts are marked by stars (*p* < 0.05).

Second, to separate the influence of audiovisual speech integration from sensory processing, delta responses were also examined in a control region of non-interest (RONI; Figure 1C). The region of non-interest was located in the occipital cortex where we did not expect higher audiovisual speech analysis to take place as visual information was identical between synchronous and asynchronous conditions within mask contrasts (RONI electrodes: PPO1h, PPO2h, PO3, POz, PO4, POO1, POO2, POO9h, O1, Oz, O2, POO10h, Ol1h, Ol2h, O9 and O10). We compared the effect of audiovisual asynchrony between the identified motor region (ROI) and the visual sensory area (RONI) to confirm that delta response modulations did not reflect signal processing only (Figure 4A). The mean differences of 2-3Hz delta power (NMA-NMS and HMA-HMS) were computed in the regions of interest and non-interest in the same time-window as previously (Figure 4B; ROI: NMA-NMS = 0.1 ± 0.09; HMA-HMS = −0.03± 0.19; RONI: NMA-NMS = 0.05 ± 0.10; HMA-HMS = 0.01 ± 0.24). A two-way repeated-measures ANOVA with the mean factors region (ROI or RONI) and mask (no-mask or head-mask) was performed to assess whether the responses of delta oscillations to asynchrony reflected multimodal speech analysis or purely signal processing taking place in sensory areas (i.e. visual occipital areas). Results revealed a significant interaction between region and mask (*F*(1, 22) = 5.75, *p* = 0.025; *η_p_^2^* = 0.21). First, Bonferroni-corrected pairwise comparisons showed that in the no-mask contrast the delta power difference NMA-NMS (but not HMA-HMS) was significantly greater in the region of interest than in the region of non-interest (respectively p = 0.025 and p = 0.572). Only in the region of interest the difference of power NMA-NMS was significantly greater than HMA-HMS (respectively p = 0.019 and p = 0.113). No main effect of mask (*F*(1, 22) = 0.25, *p* = 0.622; *η_p_^2^* = 0.21) or region (*F*(1, 22) = 2.18, *p* = 0.154; *η_p_^2^* = 0.09) was found.

**Figure 4.**
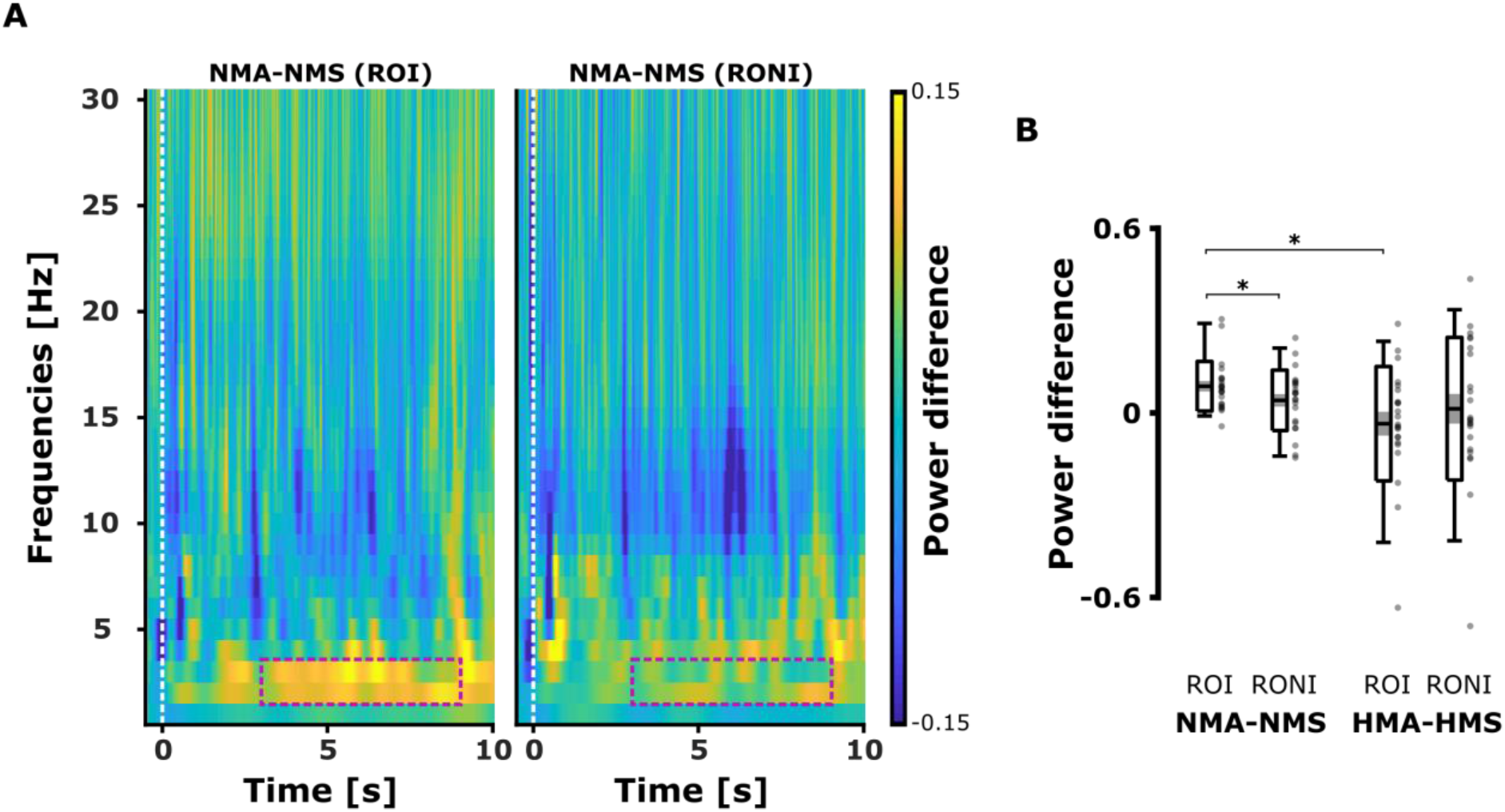
Comparisons between the motor region of interest (ROI) and the visual region of non-interest (RONI). (A) TFRs of the difference of spectrum in the no-mask contrast (NMA-NMS) in the ROI and RONI. (B) The mean differences of 2-3Hz delta power (NMA-NMS and HMA-HMS) were computed in the regions of interest and non-interest. Significant contrasts are marked by stars (*p* < 0.05).

Third, the mean power in the 4 - 8 Hz band was computed in the four conditions separately from the ROI electrodes and confirmed an increase of theta activity compared to the pre-stimulus onset baseline (NMS: 0.86 ± 0.25; NMA: 0.85 ± 0.18; HMS: 0.83 ± 0.16 and HMA: 0.81 ± 0.24). A two-way repeated-measure ANOVA revealed no significant main effect of mask (*F*(1, 22) = 2.77, *p* = 0.11; *η_p_^2^* = 0.11), synchrony (*F*(1, 22) = 0.27, *p* = 0.606; *η_p_^2^* = 0.01) or interaction between the factors mask and synchrony (*F*(1, 22) = 0.05, *p* = 0.825; *η_p_^2^* < 0.01) on theta power in the region of interest. Further, the cluster-based permutation tests revealed no significant modulation of theta power by audiovisual asynchrony in any of the mask contrasts (NMA-NMS: no significant cluster; HMA-HMS: no significant cluster; multiple comparisons are cluster-corrected). These results confirmed that audiovisual asynchrony specifically modulated delta power over the expected fronto-central region. Further, delta responses were attenuated when listeners were less able to integrate visual and auditory features, supporting the role of delta activity in the temporal integration of multimodal speech.

Next, we analysed the source localisation of the delta power modulations observed when video and audio signals were presented in asynchrony in both no-mask and head-mask contrasts. Cluster-based permutation *t*-tests between synchronous and asynchronous conditions revealed that asynchrony significantly increased delta oscillation responses when the head of the speaker was visible (NMA-NMS: *p* = 0.042; cluster statistic = 233.02) but not when it was head-masked (HMA-HMS: *p* = 0.27; cluster statistic = 38.27). The projections of the significant *t*-values on the brain’s surface showed an increase of delta power originating mainly in the left precentral region and the left inferior frontal gyrus (Figure 5A). The source results support the topographies of the delta power modulations observed at the scalp level, which revealed fronto-central differences in the no-mask contrast only (Figure 3B). Further, we tested whether delta power responses from the left motor areas correlated with the synchrony detection performance in the no-mask and head-mask contrasts (Figure 5B). Pearson correlations revealed a positive correlation between the hit rates and delta power differences in the no-mask contrast (NMA-NMS: r = 0.45; *p* = 0.031, two-tailed), but not in the head-mask contrast (HMA-HMS: r = 0.03; *p* = 0.9, two-tailed). These results confirmed that when participants were able to perceive synchrony between video and audio signals (no-mask contrast), the amplitude of delta power modulations positively correlated with accuracy. In contrast, when participants were less able to discriminate temporal alignment between visual and auditory information (head-mask contrast), left motor delta oscillations did not significantly correlate with behavioural performance.

**Figure 5.**
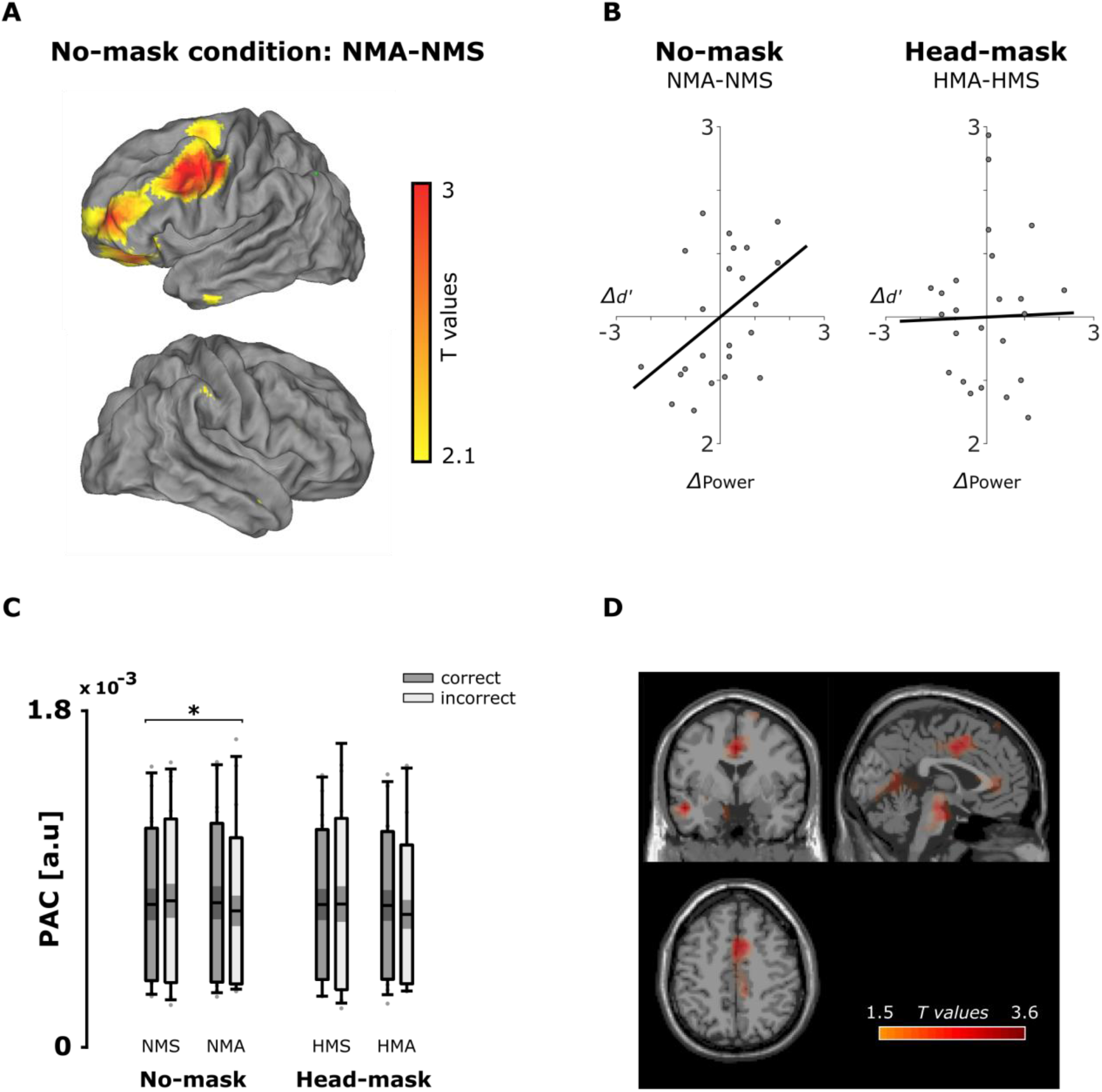
Delta oscillation responses to audiovisual asynchrony at the source level for no-mask and head-mask contrasts. (A) Contrast NMA_hit_ - NMS_hit_ projected onto the brain’s surface (significance *t*-values; cluster-corrected at alpha threshold = 0.05). The maximum voxel MNI coordinates is located left precentrally [−50 19 40] but significant activations were also found in the left inferior frontal gyrus (pars triangularis; maximum voxel MNI coordinates [−30 31 0]). No significant difference was found when the head of the speaker was masked (HMA_hit_ - HMS_hit_ contrast; not represented). (B) Scatterplots of audiovisual synchrony detection performance and delta power in the significant cluster region (left frontal-motor area). The difference of delta power in the left motor cluster (x-axis) correlated with the difference of audiovisual synchrony perception (y-axis) between synchronous and asynchronous conditions only when the face of the speaker was visible and participants could integrate video and audio onsets (no-mask contrast). (C) PAC analysis in the left frontal motor cluster. The figure represents the modulation of delta-beta PAC in a significant cluster, depending on the mask and audiovisual synchrony. Significance is indicated by an asterisk (*p* < 0.05, Bonferroni-corrected). Delta-beta PAC from the left frontal motor area was greater in the no-mask as compared to the head-mask contrast in general, but did not discriminate between hit and miss trials. Significant contrasts are marked by stars (*p* < 0.05). (D) Delta-beta PAC difference between no-mask (NMA_hit_ + NMS_hit_) and head-mask (HMA_hit_ + HMS_hit_) contrasts in the whole brain. Results revealed significant maximum differences located in the superior motor area (MNI coordinates [0 11 50]) and in the left middle temporal lobe (MNI coordinates [−50 −1 −20]).

### Delta-beta PAC reflects temporal integration in audiovisual speech perception, but is not limited to the left motor region

Finally, we assessed whether delta-beta PAC modulations in the left frontal-motor area would reflect sensitivity to audiovisual synchrony in speech. First, a three-way repeated-measure ANOVA (main factors: mask, synchrony and correctness) revealed a main effect of mask on delta-beta PAC with delta-beta phase-coupling being significantly greater in the no-mask than in the head-mask contrast (F(2,22) = 4.72; *p* = 0.041; *η_p_^2^* = 0.18; see Figure 5C). No further significant main effect or interaction were found. These results suggest that left frontal-motor delta-beta PAC relates to the temporal integration of audiovisual speech as it responded depending on whether listeners were able to match visual and auditory prosodic features (no-mask contrast) or not (head-mask contrast). Second, we investigated whether the delta-beta PAC difference between no-mask and head-mask contrasts was restricted to the left motor areas. As accuracy and synchrony did not affect delta-beta PAC in the cluster of interest, we selected only the hit trials for the delta-beta PAC analysis at the whole brain level and put synchronous and asynchronous trials together within contrasts (i.e. NMCs: NMA_hit_ + NMS_hit_; HMCs: HMA_hit_ + HMS_hit_). The cluster-based permutation tests revealed one significant positive cluster peaking in the superior motor area and in the left middle temporal lobe (although not exclusively; see Figure 5D), confirming that delta-beta PAC was significantly greater in the no-mask (NMCs) contrast as compared to the head-mask (HMCs) contrast (NMCs - HMCs: *p* = 0.043, cluster statistic = 216.69).

In summary, the EEG results mirrored the behavioural evidence as delta responses were distinctly modulated by audiovisual synchrony only when participants could view the face and visible articulators (no-mask contrast). Delta activity from the left motor region increased when visual and auditory information were misaligned (NMA), reflecting greater difficulty to match visual and auditory prosodic features as compared to the synchronous condition (NMS). Further, left frontal-motor delta responses predicted synchrony detection performance only when participants were able to properly integrate visual and auditory features (no-mask contrast), but not when they guessed (head-mask contrast). Finally, the cross-frequency coupling analysis showed that delta-beta PAC in the left frontal motor cluster of interest also increased when listeners were able to match prosodic features between modalities (no-mask contrast) as compared to when they guessed (head-mask contrast). These results suggest that delta-beta PAC in expected motor areas (although not exclusive) are sensitive to temporal integration of audiovisual speech information, and may predict whether listeners integrate visual and auditory prosodic features in asynchrony detection.

## DISCUSSION

The present study investigated the role of motor delta oscillations during the temporal integration of multimodal prosodic features in speech perception. Behavioural results showed that listeners processed both prosodic features in multimodal speech with sufficient visual information. At the brain level, the perception of audiovisual asynchrony induced an increase in delta activity in the expected left motor cortex (extending to the inferior frontal gyrus), which correlated with the participants’ sensitivity to audiovisual synchrony. In contrast, participants were less able to discriminate audiovisual synchrony when the speaker’s facial information was masked, which was characterised by an absence of delta activity response in the EEG. Finally, delta-beta PAC in the left frontal-motor areas decreased significantly when listeners could not integrate efficiently visual and auditory prosodic features in speech perception. Altogether, our results indicate that the delta time-scale provides a flexible framework to synchronise the listener’s brain activity with the temporal organization of external audiovisual speech. In this framework, the oscillatory activity can gather and realign multiple temporal representations of the visual and auditory speech features in the left motor cortex to improve dynamic signal processing.

Synchrony detection performance confirmed our first hypothesis that listeners integrate prosodic events in multimodal speech perception. This finding was expected as visual information complements auditory information and often improves speech perception (Sumby & Pollack, 1954; van Wassenhove et al., 2005). Speaker’s articulatory movements and gestures temporally aligned with acoustic prosodic cues, providing listeners with a reliable delta temporal structure of the speech signal (Biau et al., 2016; Esteve-Gibert & Guellaï, 2018; Wagner et al., 2014). Therefore, participants likely use these salient prosodic events as landmarks present in two different sensory streams to align and integrate them into a coherent multisensory speech percept. These results suggest that the temporal structure focuses the listeners’ attention within brief time-windows containing common multimodal prosodic events to facilitate their integration. This is in line with the theory of dynamic attending stating that non-random external stimulation drives periodic attention allocation towards critical events (Large & Jones, 1999). In contrast, when the speaker’s face was masked, participants could not integrate the temporal correspondence between visual and auditory prosodic anchors properly, and were less able to perceive multimodal speech. Noteworthy, the differences of performance between the no-mask and head-mask contrasts indicate that participants likely relied on complementary information conveyed by the speaker’s head, face, and fine articulatory gesture information to achieve the integration of the visual prosodic signal (Cross, Butler, & Lalor, 2015).

Further, the EEG results revealed an increase in motor delta activity in response to audiovisual asynchrony, confirming its role in temporal integration of multimodal prosodic features. Previous literature associated motor delta oscillations with the perception of rhythmic auditory inputs (Keitel et al., 2018; Morillon et al., 2019; Morillon & Schroeder, 2015). The present results extend these findings to the temporal integration of non-isochronous events that act as punctual “snap fasteners” integrating visual and auditory signals within relevant time-windows. As long as they provide the brain with a dominant temporal structure aligning multiple sensory inputs, salient prosodic features do not have to be perfectly regular to engage delta responses in the motor cortex. The present EEG results corroborate this hypothesis in three ways: First, we did not observe any different delta responses in auditory and visual cortices when audiovisual stimuli were synchronous. This would have reflected low-level feature tracking during early sensory processing (Cross, Butler & Lalor, 2015; Ghitza, 2017; Gross et al., 2013; Mai, Minett & Wang, 2016). Next, audiovisual asynchrony would likely decrease pure entrainment by making signal tracking more difficult than when different channels of the same input are processed in synchrony. Further, we found no theta activity in response to audiovisual asynchrony that would have indicated an effect driven specifically by the prosodic features’ rate (e.g., lip movements) rather than temporal integration of sensory input. Second, the difference in delta power in the left motor cortex correlated positively with performance between the synchronous and asynchronous conditions in the no-mask contrast. Moreover, the fact that performance in the synchronous and asynchronous conditions was similar when the face of the speaker was visible suggests an increase of difficulty to integrate the two temporal representations of speech signal. This extra cognitive load may be reflected by an increase of delta activity responses in the left motor cortex. Third, participants did not perceive audiovisual synchrony when the speaker’s facial information was blurred, which was reflected by weaker responses in the left motor cortex and no significant difference between synchronous and asynchronous conditions. Importantly, the responses found in the left inferior frontal gyrus align well with previous research that established a role in crossmodal information integration between gestures and speech (Park et al., 2018; Willems, Ozyürek & Hagoort, 2009; Zhao et al., 2018). Here, participants perceived information carried in the two modalities and likely integrated gestures’ kinematics with auditory envelope modulations to perform the synchrony detection task. Further investigations will need to address whether the response modulations in the left IFG were specific to gesture-speech temporal integration or could be reproduced using moving dots following gestures’ dynamics. In contrast, we found no difference of activation in additional regions associated with multimodal speech integration such as the left posterior superior temporal sulcus (Marstaller & Burianová, 2014). However, it is possible that in the present context delta oscillations did not reflect multisensory integration *per se* but temporal integration taking place in the left motor cortex and IFG.

Finally, the cross-frequency coupling analysis revealed that delta-beta coupling in the left frontal motor cortex increased when listeners perceived audiovisual (mis)alignment (no-mask contrast). This finding indicates that delta-beta PAC contributes to temporal integration of prosody as well. Potentially, delta-beta coupling may support the latter mechanisms taking place after proper temporal integration of the visual and auditory prosodic features, e.g. auditory-motor communication. Park et al. (2015) showed that the left frontal-motor areas modulated the phase of delta oscillations in the left auditory cortex by means of top-down control in speech perception. Reciprocally, delta-beta PAC in the auditory cortex respond to the modulations of rhythmic regularity in auditory speech perception (Chang, Bosnyak and Trailor, 2019). Further, Keitel et al. (2018) reported that delta-beta PAC in the left motor cortex predicted behavioural performance in speech comprehension. Future research will need to unravel whether delta-beta coupling provides a ubiquitous means of cross-regional communication to align temporally different dynamic inputs in sensory cortices (Arnal, 2012; Fujioka, Ross & Trainor, 2015; Morillon et al., 2019). For instance, Fontolan et al. (2014) reported that delta-beta coupling in the associative auditory cortex modulated the phase of gamma activity related to phonological processing in the primary auditory cortex in auditory sentence perception (Giraud & Poeppel, 2012). Alternatively, delta-beta PAC may drive the periodicity of attention to critical time-windows containing relevant accentuated speech information, which fits with the dynamic attention theory (Large & Jones, 1999).

To our knowledge, our results show for the first time how delta activity provides an interface between external dynamic stimulation and inner brain oscillations to facilitate multimodal speech perception. We propose that motor delta oscillations align together distinct representations of non-verbal and auditory prosodic features encoded separately in their respective sensory cortices. The slow time-scale of delta (1-3Hz) may also offer the brain some flexibility to create a coherent multimodal percept despite the natural delay between visual and auditory signal onsets in speech (Chandrasekaran et al., 2009). In social interactions where conditions change quickly, such a delta framework would help listeners to align the related speech streams in a bottleneck fashion to maintain a stable synchronization with the speaker’s flow (Kotz, Ravignani & Fitch, 2018). In contrast, when the temporal structure of events from two contemporary streams cannot be integrated in critical delta time-windows, they are discriminated against each other. When video and audio signal onsets were misaligned by 400ms, the alignment of the visual and auditory neural representations in the delta-phase became likely impossible, leading to the detection of asynchrony. Further investigations will need to address whether this potential mechanism exists with other time-scales present in both speech signal and endogenous oscillations. For instance, we cannot fully discard that the prosodic contour in our stimuli still contained a syllable structure embedded in it (e.g. at onsets and stress peaks). Further, lip movements and auditory envelope convey syllabic information occurring at a theta rate (4-8 Hz) providing other robust temporal information in the speech signal during face-to-face conversations (Chandrasekaran et al., 2009; Giraud & Poeppel, 2012). Therefore, delta and theta activities may actually couple to strengthen speaker-listener synchronization in social communicative interactions. Future research needs to investigate the potential role of a delta-theta coupling in speech perception.

## CONCLUSION

Our findings show that delta power and delta-beta phase-amplitude coupling in the left motor cortex reflect the temporal integration of visual and auditory prosodic events, and shaped multimodal integration in speech perception. We propose that the delta time-scale provides a reliable framework allowing endogenous activity to align multiple prosodic features conveyed in distinct sensory modalities in a common temporal organization during speech perception.

## ACKNOWLEDGMENT

This research was supported by a postdoctoral fellowship from the European Union’s Horizon 2020 research and innovation program, under the Marie Sklodowska-Curie grant agreement (No. 707727).

## RESOURCE SHARING

Consent for sharing data at the level of the individual participant was received. Data for individual participants and associated scripts will be made available after publication of the manuscript. Further information or requests should be directed to the corresponding authors.

